# Attentional prioritization enhances the accessibility of neural representations during working memory maintenance

**DOI:** 10.64898/2026.05.04.722761

**Authors:** Mingmin Zhang, Elkan Akyürek, Wouter Kruijne

## Abstract

Given the limited capacity of working memory (WM), prioritization is essential for efficient information processing. Whether prioritization acts primarily at encoding, or dynamically shapes representations during maintenance, is currently unclear. Here, we employed a delayed response task and compared prioritization conditions in which the testing order of items was either known in advance or not. Behaviorally, prioritization selectively reduced guess rates and had little effect on precision. Using multivariate pattern analysis, we decoded stimulus information from EEG voltage as an index of item specific representations, and used alpha power decoding as an index of internal attention. Prioritization did not alter decodable representations during encoding. During maintenance, however, prioritization enhanced both voltage-based decoding and alpha power-based decoding for the prioritized item. Mediation analyses further revealed an indirect association between alpha-based attentional signals and behavior via voltage-based representational strength, consistent with the idea that internal attention supports performance by maintaining prioritized representations in an accessible state.

## INTRODUCTION

Working memory (WM) temporarily stores and manipulates information, but is limited in capacity^1,2^. Whether these constraints reflect discrete slots^3–5^ or a continuous resource^6–8^, WM requires mechanisms for prioritizing information according to behavioral relevance. State-based theoretical models of WM accordingly suggest that information held in WM can occupy different priority states, with the highest priority information occupying the focus of attention and being most directly accessible for upcoming behavior^9,10^. Following these accounts, prioritization reflects the flexibility of WM^11^ and can be understood as the process by which different memory items are aligned to current or anticipated task demands.

Prioritization may act on representations in WM during their encoding and/or their maintenance. The distinction between these subprocesses of WM has been examined using various “delayed response” paradigms, in which information is encoded and subsequently maintained across a delay^12^. Pre-cues presented before stimulus onset guide prioritization during encoding and improve performance by enhancing accuracy and reducing interference^13–15^. By contrast, retro-cues presented during the delay period indicate which encoded item is most likely to be tested, thereby guiding prioritization during maintenance^15–17^ and improving memory performance for that item^18–20^. These findings demonstrate that explicit item-specific cues can guide prioritization at different stages of WM.

However, prioritization is not always triggered by a discrete cue. It can also be guided by temporal expectations about when each memory item will be required for behavior. When multiple items remain task-relevant but are expected to be used at different times, items needed sooner can be assigned higher priority than items needed later. Previous studies have shown that temporal expectations facilitate access to WM contents at their expected time of use and are accompanied by temporally specific changes in alpha band activity during maintenance^21,22^. Specifically, when the order of two memory tests was known in advance, the item tested first showed more robust decoding than the item tested second^23^. Recent work similarly found that both items were initially decodable following stimulus presentation, but sustained voltage- and alpha power-based decoding throughout the delay was primarily observed for the item tested first^24^. However, because previous studies have typically compared first- and second-tested items only when test order is known, without a condition where order is unpredictable, it remains unclear when prioritization guided by anticipated test order emerges.

One possibility is that such prioritization emerges during encoding. Advance information about test order may establish an anticipatory attentional set before stimulus onset, allowing attention to be oriented toward the item expected to be tested first as it is encoded^15,25^. A second possibility is that prioritization emerges during maintenance. Both items may initially be encoded without differential priority, after which anticipated test order guides internal attention toward the first tested item during the delay. Temporal expectations could thereby dynamically adjust priority during the delay^21,22,24,26^. A third possibility is that prioritization unfolds across both stages, with selection beginning during encoding and subsequently strengthening during maintenance to make the first tested item more accessible for upcoming readout^9,23^.

To determine whether prioritization guided by anticipated test order emerges during encoding, during maintenance, or across both stages, we designed an EEG experiment that manipulated priority information. Participants remembered two oriented gratings, and after a delay, compared each item to a probe in turn (Figure 1A). In fixed-order blocks, participants knew in advance which item would be tested first, making item specific temporal priority available before encoding. In random-order blocks, the test sequence was unpredictable, such that neither item was favored by advance order information before the first test. During maintenance, impulse stimuli were presented before each probe to allow neural readout of the memoranda^23^. To examine how prioritization shapes memory representations during encoding and maintenance, we separately decoded item-specific information from voltage and alpha power. We decoded from alpha power specifically to test the degree to which priority-dependent changes were associated with internal attentional selection^24^.

**Figure 1.**
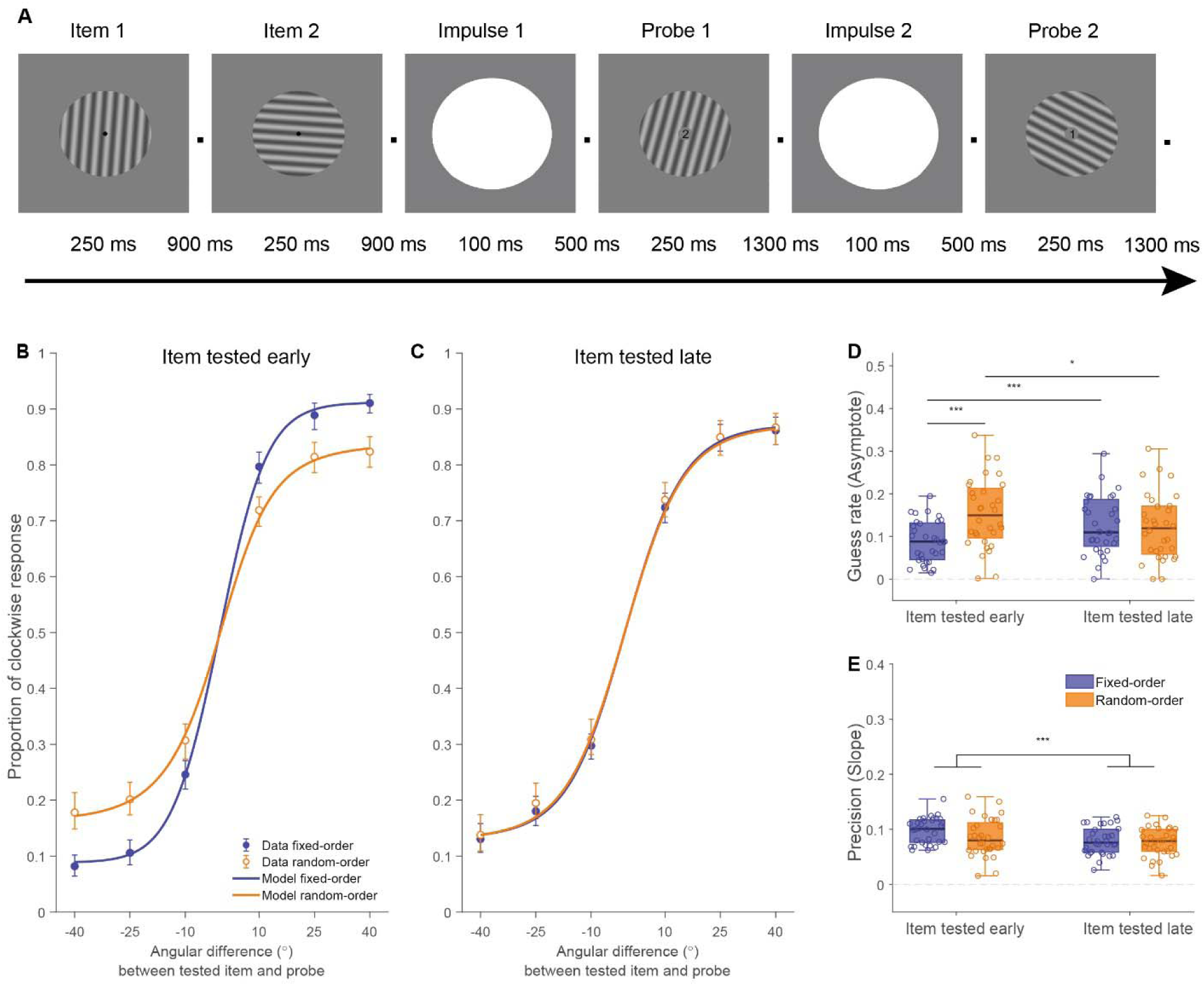
Task Procedure and WM task performance in fixed- and random-order conditions across all probes. (A) The sequence of events within a trial is shown. Two memory items were presented sequentially, followed after blank delay intervals by Impulse 1, Probe 1, Impulse 2, and Probe 2. (B&C) Probability of clockwise responses as a function of the target-probe angular difference for the tested-early and tested-late items. Lines show model fits. Error bars denote 95% within-participant bootstrap confidence intervals. (D) Guess rate and (E) precision for early and late items in the fixed-order and random-order conditions. Asterisks indicate significant differences: ** p < 0.01, *** p < 0.001.

We furthermore implemented impulse perturbation, because while item-specific information can be decoded from stimulus-evoked neural activity during encoding^23,27^, delay period activity is often dynamic or intermittent rather than continuously sustained, such that maintained information may not remain consistently decodable from ongoing neural activity^28,29^. These observations have motivated activity-silent or -quiescent accounts, according to which WM information can be retained in latent changes in network state, potentially supported by short-term synaptic plasticity, without continuous content-specific neural activity^10,23,30–32^. Here we presented a task-irrelevant visual impulse during maintenance to perturb the memory network and evoke a response that reflects its current state, to reveal item-specific and potentially priority-dependent information that is not evident in ongoing neural activity^23,33,34^.

## RESULTS

### Behavioral Results

Figure 1B and C present the behavioral results for the two items and prioritization condition. The analysis of guess rate, derived from these model fits, revealed a main effect of prioritization condition (Figure 1D; *F*(1, 31) = 12.427, *p* = 0.001, *η_p_*^2^ = 0.286), but no effect of tested item (*F*(1, 31) = 0.270, *p* = 0.607, *η_p_*^2^ = 0.009). Their interaction was significant (*F*(1, 31) = 13.077, *p* = 0.001, *η_p_*^2^ = 0.297). Post-hoc analyses revealed that guess rates for the item tested early were significantly lower in the fixed-order condition (*M* = 0.089, *SD* = 0.049) than in the random-order condition (*M* = 0.153, *SD* = 0.081; *t*(31) = -5.584, *p* < 0.001, *d* = -0.987, 95% CI [-0.089, -0.041]). For the item tested late, no significant difference was observed between the fixed-order condition (*M* = 0.126, *SD* = 0.068) and random-order condition (*M* = 0.122, *SD* = 0.077; *t*(31) = 0.326, *p* = 0.746). When contrasting the items tested early and late, guess rates were significantly lower for the item tested early in the fixed-order condition (*t*(31) = -4.913, *p* < 0.001, *d* = −0.869, 95% CI [−0.054, −0.022]), whereas they were significantly higher for the item tested early in the random-order condition (*t* (31) = 2.258, *p* = 0.031, *d* = 0.399, 95% CI [0.003, 0.060]).

For precision (slope; Figure 1E), a significant main effect of tested item was found (*F*(1, 31) = 21.705, *p* < 0.001, *η_p_*^2^ = 0.412), whereas neither the main effect of prioritization condition (*F*(1, 31) = 3.239, *p* = 0.082, nor the interaction (*F*(1, 31) = 1.371, *p* = 0.251) was significant. This main effect reflected higher precision for the item tested early (*M* = 0.092, SD = 0.022) than for the item tested late (*M* = 0.078, SD = 0.020; *t*(31) = -4.659, *p* < 0.001, *d* = 0.824, 95% CI [0.008, 0.020]). The descriptive statistics were as follows: item tested early in the fixed-order condition (*M* = 0.098, *SD* = 0.024), item tested early in the random-order condition (*M* = 0.085, *SD* = 0.034), item tested late in the fixed-order condition (*M* = 0. 079, *SD* = 0.025), and item tested late in the random-order condition (*M* = 0.076, *SD* = 0.026).

These results suggest that prioritization primarily affected guess rates rather than precision, with reduced guessing for the item tested early in the fixed-order condition. For the item tested late, which was always predictable and thus could be prioritized, guess rate was not different between the conditions.

### Voltage Decoding Results

#### Encoding phase

We first examined decoding of the item tested early from EEG voltages during the encoding phase. As shown in Figure 2A and Figure 2B, following the presentation of Item 1, the item tested early could be successfully decoded in both the fixed-order (82-714 ms, p < 0.001) and random-order conditions (58-586 ms, p < 0.001). Late within these clusters, we also observed that decoding accuracy was significantly higher in the fixed-order condition than the random-order condition (474-554 ms, p = 0.017). Following the presentation of Item 2, successful decoding of the item tested early was again observed in both fixed-order (74-594 ms, p < 0.001) and random-order condition (98-618 ms, p < 0.001), with no significant differences between conditions.

**Figure 2.**
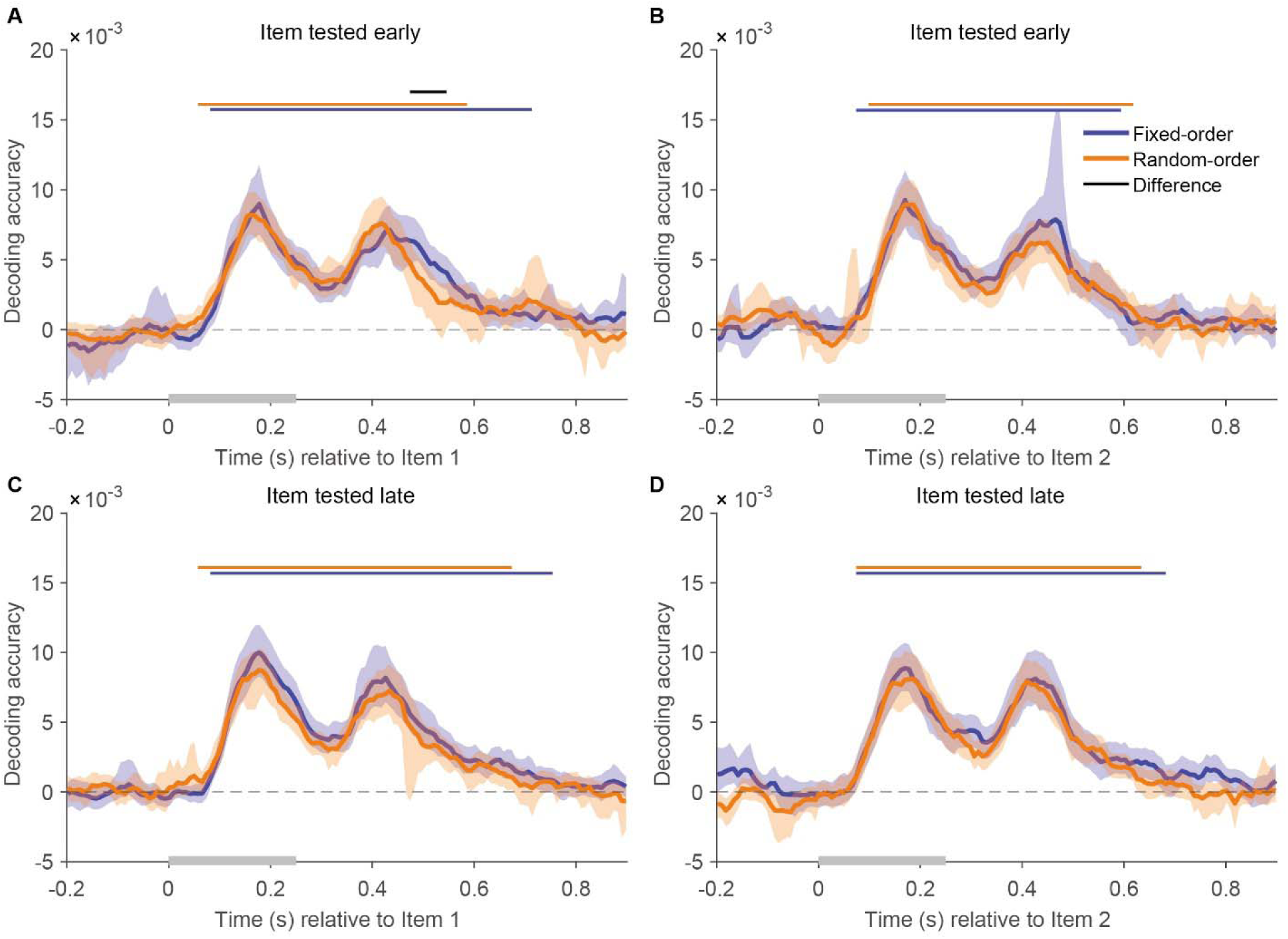
Voltage decoding during encoding. Time-resolved voltage decoding accuracy is shown separately for the item tested early (A-B) and the item tested late (C-D), time-locked to Item 1 (A, C) and Item 2 (B, D). Shaded bands denote 95% confidence intervals. Colored horizontal bars indicate clusters with decoding significantly above chance in that condition (cluster-based permutation test). The black bar (in A) indicates a significant difference between conditions. Gray bars along the x-axis indicate the item presentation period.

Decoding of the item tested late is shown in Figure 2C and Figure 2D. Following the presentation of Item 1, the item tested late could be successfully decoded in both the fixed-order (82-754 ms, p < 0.001) and random-order conditions (58-674 ms, p < 0.001). No significant differences were observed between these conditions. After the presentation of Item 2, successful decoding of the item tested late was again observed in both fixed-order (74-682 ms, p < 0.001) and random-order condition (74-634 ms, p < 0.001), with no significant differences between them.

Taken together, these findings show that both items could be reliably decoded from EEG voltages during encoding. Although decoding of the item tested early was briefly higher in the fixed-order than in the random-order condition following Item 1, supplementary within-condition contrasts comparing early-versus late-tested items revealed no significant differences in either encoding epoch, in either prioritization condition (Supplementary Figure S2). Thus, the encoding data provide little evidence that future priority status reliably modulated voltage decoding during encoding.

#### Maintenance phase

We next examined voltage decoding results during the maintenance phase, in response to the impulse stimuli. Following Impulse 1, reliable decoding of the item tested early (Figure 3A) was observed in the fixed-order condition (82-426 ms, *p* = 0.011). Critically, direct comparisons revealed significantly greater decoding accuracy in the fixed-order condition than in the random-order condition (122-410 ms, *p* = 0.001). Following Impulse 2 (Figure 3B), when this item had already been tested and was no longer relevant, we observed successful decoding in both the fixed-order (266-354 ms, *p* = 0.024) and random-order (138-410 ms, *p* < 0.001) conditions. No significant differences were found between these conditions.

**Figure 3.**
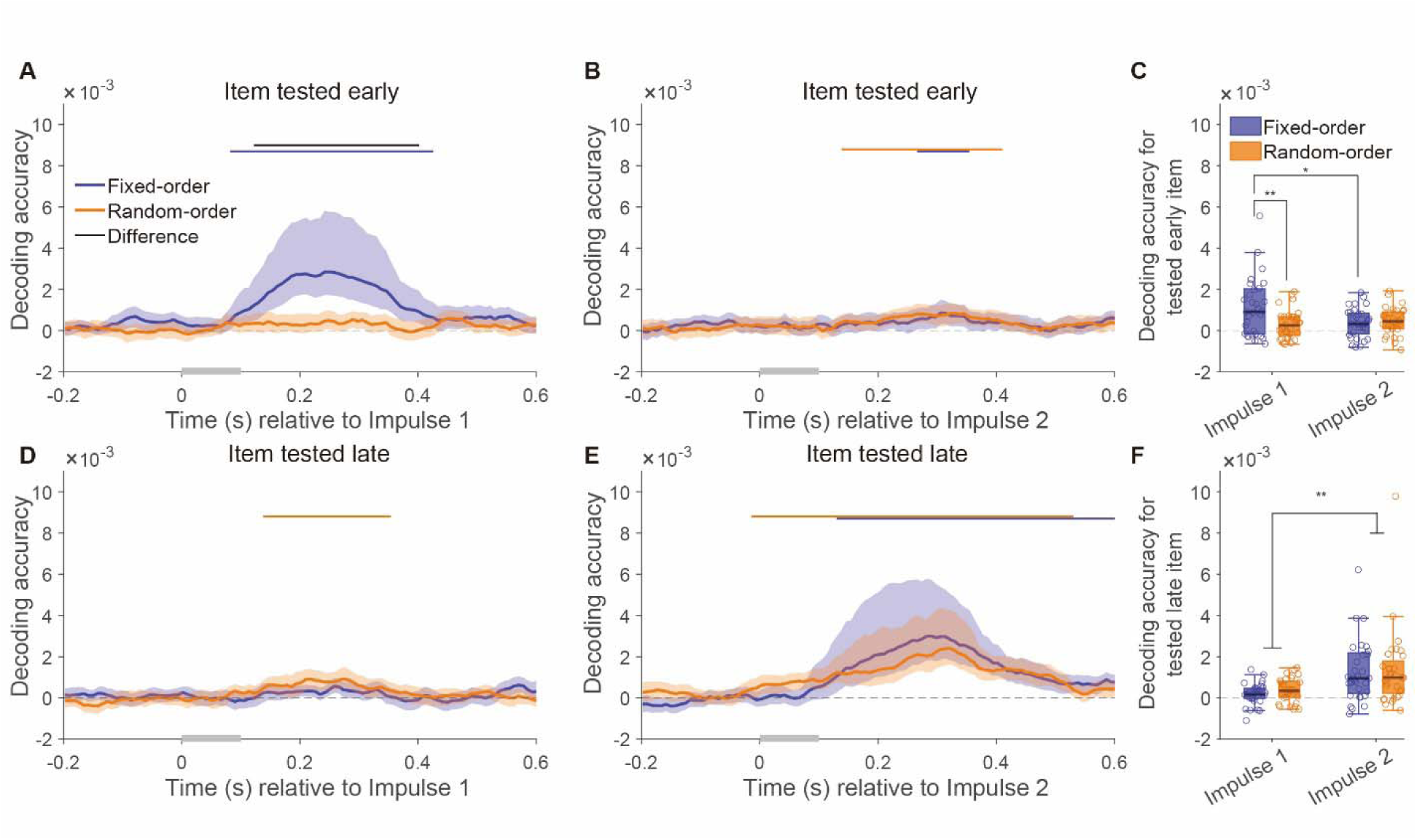
Time-resolved voltage decoding following impulses. Voltage decoding accuracy is shown time-locked to Impulse 1 (A, D) and Impulse 2 (B, E) for the item tested early (A-C) and the item tested late (D-F). Panels C and F show decoding accuracy averaged over the impulse window (100-600 ms). Asterisks indicate significant pairwise differences: * p < 0.05; ** p < 0.01.

We next conducted a two-way repeated-measures ANOVA on decoding accuracy for the item tested early, averaged over 100-600 ms after each impulse (Figure 3C). The analysis again revealed that the main effect of prioritization condition was significant (*F*(1, 31) = 5.194, *p* = 0.030, *η_p_*^2^ = 0.144), while the main effect of impulse epoch was not (*F*(1, 31) = 3.808, *p* = 0.060, *η_p_*^2^ = 0.109). Importantly, their interaction was significant (*F*(1, 31) = 10.751, *p* = 0.003, *η_p_*^2^ = 0.258), and post-hoc analyses revealed the source of this interaction: For Impulse 1, decoding accuracy in the fixed-order condition (*M* = 1.494 × 10^-3^, *SD* = 2.649 × 10^-3^) was significantly higher than in the random-order condition (*M* = 0.301 × 10^-3^, *SD* = 0.679 × 10^-3^; *t*(31) = 2.926, *p* = 0.006, *d* = 0.517, 95% CI [0.361 × 10^-3^, 2.020 × 10^-3^]). For Impulse 2, decoding accuracy in the fixed-order condition (*M* = 0.383 × 10^-3^, *SD* = 0.700 × 10^-3^) did not significantly differ from that in the random-order condition (*M* = 0.450 × 10^-3^, *SD* = 0.649 × 10^-3^; *t*(31) = -0.381, *p* = 0.706). In line with this pattern, we additionally found that in the fixed-order condition, decoding accuracy for Impulse 1 was significantly higher than for Impulse 2 (*t*(31) = 2.686, *p* = 0.012, *d* = 0.475, 95% CI [0.267 × 10^-3^, 1.953 × 10^-3^]), whereas in the random-order condition, decoding accuracy following Impulse 1 did not significantly differ from that following Impulse 2 (*t*(31) = -0.956, *p* = 0.347).

Focusing on the item tested late, following Impulse 1 (Figure 3D), reliable decoding was observed only in the random-order condition (138-354 ms, *p* = 0.002). In the fixed-order condition, where the item tested late was not yet relevant, decoding was qualitatively elevated following the impulse, but no significant between condition differences were found. Following Impulse 2 (Figure 3E), when the item tested late was about to be probed, decoding was robust in both the fixed-order (130-600 ms, *p* < 0.001) and random-order (-14-530 ms, *p* = 0.005) conditions, with no significant differences between conditions.

The two-way repeated-measures ANOVA’s on mean decoding accuracy for the item tested late (Figure 3F) revealed that the main effect of impulse epoch was significant (*F*(1, 31) = 9.991, *p* = 0.004, *η_p_*^2^ = 0.244). However, we found no main effect of prioritization condition for the item tested late (*F*(1, 31) = 0.339, *p* = 0.564, *η_p_*^2^ = 0.011), and no significant interaction effect (*F*(1, 31) = 3.943, *p* = 0.056, *η_p_*^2^ = 0.113). Consistent with the significant main effect of impulse epoch, post-hoc tests confirmed the decoding accuracy was significantly higher for Impulse 2 (*M* = 1.485 × 10^-3^, *SD* = 2.247 × 10^-3^), compared to Impulse 1 (*M* = 0.264 × 10^-3^, *SD* = 0.373 × 10^-3^; *t*(31) = 3.161, *p* = 0.004, *d* = 0.559, 95% CI [0.429 × 10^-3^, 1.993 × 10^-3^]).

In summary, for the item tested early, decoding accuracy following Impulse 1 was significantly higher in the fixed-order condition than in the random-order condition. This difference between conditions subsequently disappeared after this item had been tested, with this item showing only marginally above chance decoding following Impulse 2, in both prioritization conditions. Conversely, the item tested late did not show any significant between condition differences at either time point; instead, decoding accuracy for this item was low in response to Impulse 1 but was significantly increased in both conditions following Impulse 2, which is right before it got probed. These results show that the representational strength, indexed by voltage-based decoding, depends on the item’s prioritization status. In the fixed-order condition, items that were about to be tested were decoded better than irrelevant items. In the random-order condition, this only held for the item tested late, as the item tested early would not yet be known to the participant during the first delay.

### Alpha Power Decoding Results

For alpha power decoding of the item tested early, we found seemingly sustained decoding above chance in the fixed-order condition throughout the entire epoch. That is, we found a cluster of above chance decoding in the fixed-order condition, that seemed to be in place already before the onset of Impulse 1 (Figure 4A; -200-270 ms, *p* = 0.010), shortly thereafter followed by a cluster that lasted until the end of the epoch (398-600 ms, *p* < 0.001). Conversely, in the random-order condition there was no evidence that decoding scores were significantly above chance. Direct comparisons revealed significantly higher decoding accuracy in the fixed-order condition than in the random-order condition (412-600 ms, *p* < 0.001). During the second delay (Figure 4B), decoding of the item tested early was not significant in the fixed-order condition, but significant decoding of this item was found in the random-order condition, early in the epoch, before the impulse stimulus (-200-26 ms, *p* = 0.048). No significant difference was observed between the fixed-order and random-order conditions.

**Figure 4.**
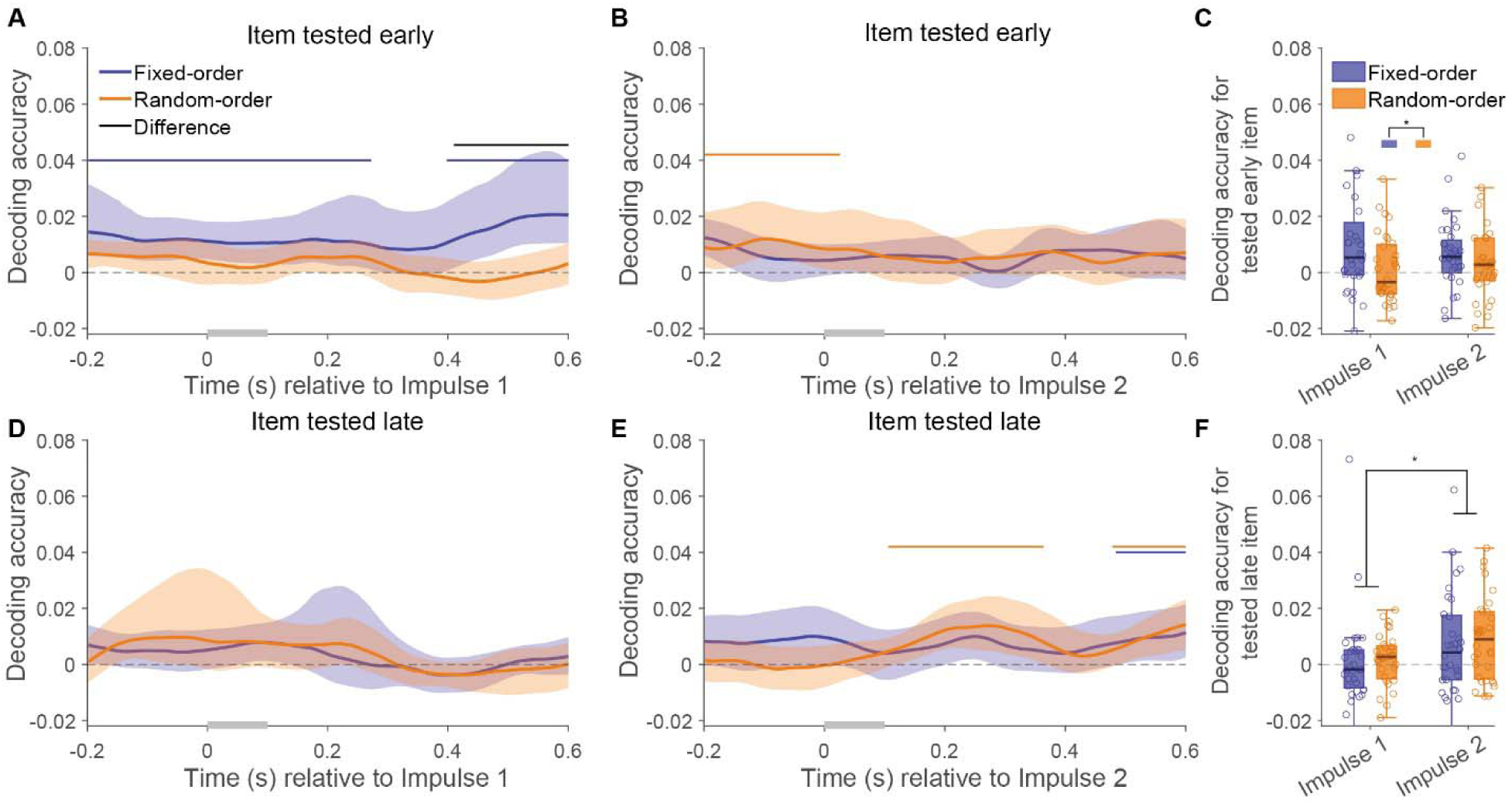
Time-resolved alpha power decoding following impulses. Alpha power decoding is shown time-locked to Impulse 1 (A, D) and Impulse 2 (B, E) for the item tested early (A-C) and the item tested late (D-F). Panels C and F show decoding accuracy averaged over the impulse window (100-600 ms).

The two-way repeated-measure ANOVA results for the item tested early (Figure 4C) revealed no significant main effect of impulse epoch (*F*(1, 31) = 0.622; *p* = 0.436; *η_p_*^2^ = 0.020). The main effect of prioritization condition was significant (*F*(1, 31) = 5.056; *p* = 0.032; *η_p_*^2^ = 0.140). Post-hoc tests confirmed that decoding accuracy was significantly higher in the fixed-order condition (*M* = 9.216 × 10^-3^, *SD* = 17.507 × 10^-3^) than in the random-order condition (*M* = 3.336 × 10^-3^, *SD* = 11.213 × 10^-3^; *t*(31) = 2.249, *p* = 0.032, *d* = 0.398, 95% CI [0.547 × 10^-3^, 11.212 × 10^-3^]). The interaction was not significant (*F*(1, 31) = 2.935; *p* = 0.097; *η_p_*^2^ = 0.086).

Following Impulse 1 (Figure 4D), decoding of the item tested late was not significant in either prioritization condition, and no reliable difference was observed between conditions. In contrast, following Impulse 2 (Figure 4E), reliable decoding was observed in both the fixed-order (486-600 ms, *p* < 0.001) and random-order (116-364 ms, *p* = 0.027; 478-600 ms, *p* < 0.001) conditions. No significant difference was found between the fixed-order and random-order conditions.

The two-way repeated-measures ANOVA results for the item tested late (Figure 4F) revealed a significant main effect of Impulse epoch (*F*(1, 31) = 6.136; *p* = 0.019; *η_p_*^2^ = 0.165), with higher decoding accuracy following Impulse 2 (*M* = 8.049 × 10^-3^, *SD* = 15.030 × 10^-3^) than Impulse 1 (*M* = 1.040 × 10^-3^, *SD* = 10.435 × 10^-3^; *t*(31) = 2.477, *p* = 0.019, *d* = 0.438, 95% CI [1.238 × 10^-3^, 12.780 × 10^-3^]). The main effect of prioritization condition was not significant (*F*(1, 31) = 0.549; *p* = 0.464; *η_p_*^2^ = 0.017). The interaction was also not significant (*F*(1, 31) = 0.131; *p* = 0.720; *η_p_*^2^ = 0.004).

Taken together, the analysis of alpha power during maintenance qualitatively match the voltage-based analysis, in terms of the main effects of prioritization condition and impulse epoch. The item tested early revealed a significant main effect of prioritization condition, with decoding accuracy being significantly higher in the fixed-order than in the random-order condition. Conversely, the item tested late showed a significant main effect of Impulse epoch, characterized by higher decoding accuracy following Impulse 2 compared to Impulse 1, with no significant differences found between the fixed-order and random-order conditions.

### Regression And Mediation Results

The behavioral effects of prioritization in different conditions were mainly found in guess rates. Neural effects of prioritization, indexed by decoding scores in the maintenance epochs preceding the item to be probed, seemed to be reflected in elevated voltage decoding in response to the impulse stimulus, and more sustained alpha power decoding throughout the delay, (Figure 3A, E; Figure 4A, E), particularly in the fixed-order condition (see also Figure 3C, F and Figure 4C, F). We therefore next assessed whether these neural decoding measures directly related to behavioral effects. To this end, we constructed per-participant measures of these decoding accuracies by averaging voltage decoding across 100-400ms post-impulse and alpha decoding across the entire epoch (-200-600ms) Then, we constructed regression models predicting behavior from these decoding scores, separately per prioritization condition and tested item.

As shown in Table 1 and Figure S3 A-B, for the fixed-order condition, the univariate Voltage Model and Alpha Model showed that both voltage decoding and alpha power decoding were negatively associated with guess rates. The Voltage Model explained more variance (see R^2^) than the Alpha Model. The Combined Model explained the most variance, yet its AIC was higher than that of the Voltage model. For the random-order condition (Table 1 and Figure S3 C-D), the Voltage Model again showed a negative association between voltage decoding and guess rate, and explained more variance than the Alpha Model which showed a similar negative relationship between alpha power decoding and guess rate. The Combined Model explained slightly more variance than the Voltage Model, but had a higher AIC. These results suggest that voltage decoding was more closely related to WM performance than alpha power decoding, with alpha power decoding providing little additional explanatory value beyond voltage decoding.

**Table 1:** Linear regression models predicting guess rate from decoding measures.

| Predictors | *β* | SE | *R*<sup>2</sup> | AIC |
|---|---|---|---|---|
| **Fixed-order condition** |  |  |  |  |
| Voltage | -6.626 | 2.333 | 0.212 | -97.465 |
| Alpha | -1.428 | 0.603 | 0.157 | -95.331 |
| Voltage | -5.019 | 2.691 |  |  |
| + Alpha | -0.791 | 0.673 | 0.248 | -96.956 |
| **Random-order condition** |  |  |  |  |
| Voltage | -19.541 | 6.767 | 0.217 | -83.846 |
| Alpha | -4.746 | 1.438 | 0.266 | -85.916 |
| Voltage | -13.847 | 6.589 |  |  |
| + Alpha | -3.728 | 1.446 | 0.363 | -88.452 |

Note. *β* = unstandardized regression coefficient; SE = standard error; *R*<sup>2</sup> = coefficient of determination; AIC = Akaike information criterion.

However, alpha power decoding may have an indirect relationship with WM performance, a possibility we further examined using mediation analysis. We next examined the relationship between alpha power decoding and voltage decoding by fitting linear regression models with voltage decoding as the dependent variable and alpha power decoding as the predictor. Because alpha decoding of prioritized items seemed to be sustained, whereas voltage-based decoding seemed to be evoked by the impulse, we conducted a time-resolved analysis, using alpha decoding scores across the epoch to predict voltage-based decoding evoked by the impulse (Figure S3 E). These analyses revealed early and late clusters of significant positive associations in the fixed-order condition, from −58 to 340 ms and from 586 to 600 ms. In the random-order condition, only a later cluster was found from 138 to 302 ms.

Based on these findings, we hypothesized that perhaps alpha decoding throughout the delay may reflect internal attention involved in prioritization, which could enhance the accessibility of the memory representation indexed by impulse evoked voltage decoding. Therefore, we conducted separate mediation analyses using alpha decoding from the pre-impulse (−200–0 ms) and post-impulse (100–400 ms) windows as the predictor, voltage decoding averaged over 100–400 ms post-impulse as the mediator, and guess rate as the behavioral outcome. These mediation analyses (Figure 5A) tested whether the relationship between alpha power decoding and lower guess rates was mediated by voltage decoding. For this analysis, we first averaged the epochs surrounding Impulse 1 and 2 and then analyzed these separately.

**Figure 5.**
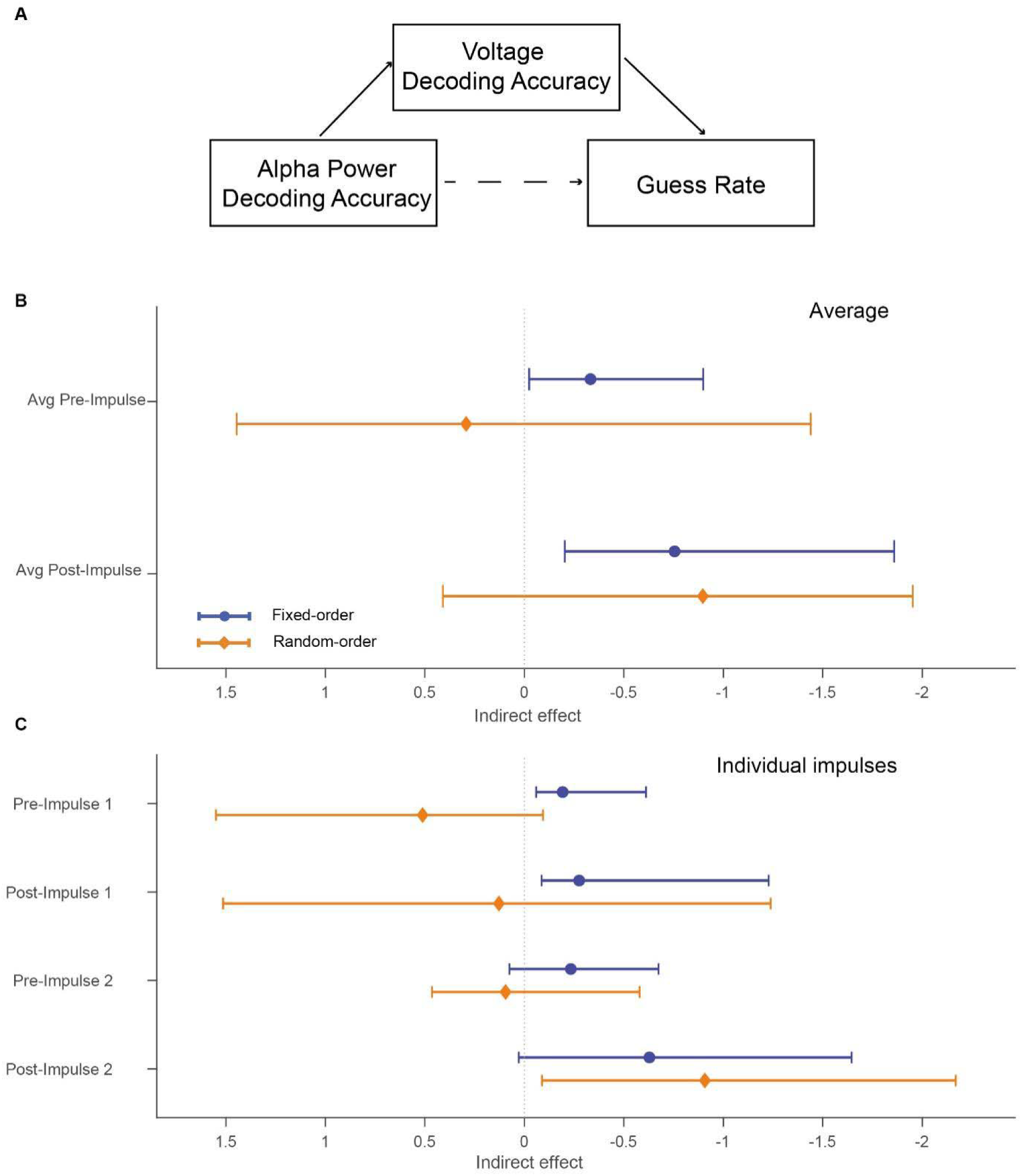
Mediation analysis of the relationship between alpha power decoding, voltage decoding, and guess rate. (A) Mediation model. (B) Indirect effects averaged across Impulses 1 and 2. (C) Indirect effects shown separately for each impulse. Note that decoding accuracies and defined with respect to the to-be-tested item only. Pre- and Post-Impulse refer to the time windows used for alpha power decoding.

When Impulses 1 and 2 were averaged (Figure 5B), we found significant support for the proposed mediation model in the fixed-order condition, as we found significant indirect effects for both pre-impulse alpha decoding (indirect effect = − 0.332, 95% CI [−0.899, −0.025]) and post-impulse alpha decoding (indirect effect = −0.755, 95% CI [−1.859, −0.204]). Neither indirect effect was significant in the random-order condition, but qualitatively there was much more evidence for mediation considering post-impulse alpha decoding than for pre-impulse alpha decoding. The impulse specific analyses (Figure 5C) showed significant indirect effects for Impulse 1 in the fixed-order condition during both pre-impulse alpha decoding (indirect effect = −0.192, 95% CI [−0.612, −0.059]) and post-impulse alpha decoding (indirect effect = −0.276, 95% CI [−1.228, −0.087]). For Impulse 2 in the fixed-order condition, the indirect effect was not significant during pre-impulse alpha decoding (indirect effect = −0.233, 95% CI [−0.674, 0.075]), or during post-impulse alpha decoding (indirect effect = −0.629, 95% CI [−1.645, 0.028]), but note that the 95% CI in the latter only narrowly crossed zero. In the random-order condition, neither indirect effect was significant for Impulse 1, neither pre-nor post-impulse (indirect effect = 0.512, 95% CI [−0.094, 1.551]; indirect effect = 0.128, 95% CI [−1.238, 1.515]). For Impulse 2, a significant indirect effect was observed during post-impulse alpha decoding (indirect effect = − 0.907, 95% CI [−2.169, −0.088]), but not during pre-impulse alpha decoding (indirect effect = 0.095, 95% CI [−0.579, 0.464]). The results suggest that whenever sufficient information is available to identify which item should be prioritized, internal attention may increase the accessibility of the representation of the prioritized item, and thereby improve WM performance.

## DISCUSSION

In this study, we first examined how prioritization in WM unfolds over time. Our results indicate that prioritization was expressed primarily during maintenance rather than encoding. Advance test-order information produced little consistent modulation of item-specific voltage decoding during encoding, whereas priority-dependent differences were found during maintenance in both voltage and alpha power decoding. We also examined how prioritization affected WM performance. Behaviorally, advance information about test order selectively reduced guess rates and had little effect on precision. Mediation analyses further suggested that internal attention may reduce guessing by maintaining prioritized representations in a state that is more readily accessible for readout during maintenance.

Behaviorally, we found that prioritization primarily affected guess rates and had little effect on precision. This dissociation suggests that prioritization increased the likelihood that the relevant representation could be accessed and used at test, rather than improving its fidelity when successfully retrieved^35,36^. At the neural level, there was little evidence that advance test-order information modulated item specific voltage decoding during encoding. Both items could reliably be decoded in the fixed-and random-order conditions. Although decoding of the item tested early was briefly higher in the fixed-order condition following Item 1, this difference did not recur following Item 2 and was not supported by direct comparisons between items tested early and late within either condition. These behavioral and neural findings differ from Wolff et al. ^23^, who reported a benefit in precision and, under simultaneous presentation, sustained decoding of the prioritized item while decoding of the non-prioritized item rapidly declined in the encoding phase. One possible explanation for the difference in neural encoding results concerns the mode of stimulus presentation. Sequential presentation may reduce direct competition, and therefore prevent encoding trade-offs between items, as each item is encoded within a separate temporal epoch^37,38^.

By contrast, we found that priority-related differences were clearly expressed during maintenance. During the first delay, the item tested early showed stronger item specific voltage decoding in the fixed-order condition than in the random-order condition, as the participants only knew which item would be tested next in the former condition. Because both items still had to be retained and tested, this difference is unlikely to reflect whether an item remained in memory. Rather, advance test-order information appears to have increased the accessibility of the next relevant representation for the forthcoming test. Similarly, the absence of an equivalent enhancement in the random-order condition does not establish that accessibility was evenly distributed across the two items; it shows only that advance information did not identify either item as the next target. After the first probe, the remaining item became the next target in both conditions. Its decoding increased before the second probe, and no difference between the fixed-and random-order conditions was observed at this stage. Thus, temporal priority was updated as the identity of the next relevant item changed, rather than being assigned during encoding and preserved unchanged throughout the trial. This interpretation is consistent with evidence that temporal expectations prepare maintained information for its expected time of use and that shifts in behavioral priority change how strongly a memory item contributes to an upcoming decision^21,22,27^. The continued decoding of the item tested early after its response further indicates that losing priority did not immediately eliminate its representation; prioritization changed its accessibility for upcoming use rather than simply determining whether it remained in memory^10^.

Within WM, prioritization may be achieved by shifting internal attention toward one representation while leaving others outside the focus of attention^39^. Internal attention has been proposed as a core mechanism for maintaining relevant items in a prioritized state^9,40^, and alpha band activity has been linked to the focus of internal attention^41,42^. Against this background, we asked whether the maintenance related prioritization effect observed in our behavioral and voltage decoding results was accompanied by a distinct attentional state. Our alpha power decoding results suggest that internal attention was most clearly engaged when priority had already been established in advance. For the item tested early, this was expressed most strongly in the fixed-order condition during the first maintenance phase. Time-resolved cluster analyses of alpha power during maintenance suggested that internal attention actively contributes to the maintenance of prioritized information during the delay, a finding that aligns with the observations of Weng et al^24^. Interestingly, in the random-order condition, the previously tested item was still decodable in pre-impulse alpha band before Impulse 2, even at the end of the response delay. This suggests that it takes a moment for attention to fully move from the old item to the new one.

Our neural decoding measures were furthermore related to behavioral performance. The regression analyses suggested that voltage decoding was more closely related to guess rate than alpha power decoding. Meanwhile, alpha power decoding and voltage decoding were also correlated. Given that alpha power decoding may index attentional selection and voltage decoding may index the accessibility of item specific information^23,41^, these findings suggest that attentional selection and representational accessibility may be related, while the representational accessibility is more closely associated with subsequent memory performance. When decoding measures were averaged across the two impulses, an indirect association between alpha power decoding and guess rate through voltage decoding was observed in the fixed-order condition, but not in the random-order condition. This pattern may reflect when participants had sufficient information to prioritize the item relevant to the upcoming test. In the fixed-order condition, the upcoming test item could be identified at both impulses, whereas in the random-order condition, it became predictable only at impulse 2, after the first item had been tested. Consistent with this account, the indirect association was observed at Impulse 1 in the fixed-order condition but not in the random-order condition, whereas at Impulse 2, it was observed in both conditions. At Impulse 2, the indirect association was observed during the early post-impulse interval in both conditions, suggesting that this relationship was expressed following the impulse when the upcoming test item could be identified. The pre- and post-impulse analyses further examined whether this relationship was associated with alpha power decoding before or after the visual impulse. Together, these findings suggest that advance knowledge of the upcoming test item may facilitate attentional prioritization, which is associated with the accessibility of task-relevant representations and subsequent WM performance.

## Conclusion

In conclusion, our findings indicate that prioritization based on advance test-order information operates primarily during WM maintenance. We found little consistent evidence that prioritization altered item specific voltage decoding during encoding. During maintenance, however, prioritization modulated both voltage and alpha power decoding, and these effects changed as the item needed next changed over the course of the trial. Behaviorally, prioritization selectively reduced estimated guess rates rather than precision. The mediation results are consistent with the possibility that prioritization improves performance by recruiting internal attention to keep the item needed next in a more accessible state. Overall, prioritization appears to support WM by dynamically adjusting the accessibility of maintained information according to upcoming behavioral demands.

## Limitations of the study

Several limitations should be noted. First, the present study included only two sequentially encoded items, which may limit the generalizability of the observed prioritization effects to WM situations involving larger set sizes. Second, although the task was intentionally demanding, participants were screened based on their performance during practice, which may have introduced sampling bias and restricted the broad applicability of the findings. Third, because the two memory items were always different within the same trial, their neural representations may have interfered with one another and therefore cannot be considered fully independent. As a result, decoding of one item may partly reflect the representation of the other, which could inflate decoding estimates. However, our analyses focused on condition-dependent changes across the delay period, so that this dependency could not account for the observed effects. Future work should investigate whether this mechanism persists under higher memory loads and extends to broader populations beyond high-performing individuals.

## RESOURCE AVAILABILITY

### Lead contact

Requests for further information and resources should be directed to and will be fulfilled by the lead contact, Mingmin Zhang.

### Materials availability

This study did not generate new unique reagents.

### Data and code availability

- Filtered and epoched EEG data and the associated behavioral data have been deposited at DataverseNL as listed in the <u>key resources table</u> and are publicly available as of the date of publication.
- All original code has been deposited at DataverseNL and is publicly available at <u>key resources table</u> as of the date of publication.
- Any additional information required to reanalyze the data reported in this paper is available from the lead contact upon request.

## ACKNOWLEDGMENTS

We thank Dionysia Kontostavlaki and Maria Ogircin for their assistance with EEG data collection, as well as the Center for Information Technology of the University of Groningen for their support and for providing access to the Hábrók high performance computing cluster. This work was supported by a grant from the China Scholarship Council (CSC202206140033 to M.Z.).

## AUTHOR CONTRIBUTIONS

Conceptualization, M.Z, E.A. and W.K.; investigation, M.Z.; formal analysis, M.Z.; visualization, M.Z. and W.K.; writing – original draft, M.Z.; writing – review & editing, M.Z, E.A. and W.K.; supervision, E.A. and W.K.

## DECLARATION OF GENERATIVE AI AND AI-ASSISTED TECHNOLOGIES IN THE WRITING PROCESS

During the preparation of this work, the first author used ChatGPT in order to improve the language of the manuscript. After using this tool or service, all authors reviewed and edited the content as needed and take full responsibility for the content of the publication.

## SUPPLEMENTAL INFORMATION

To test whether encoding activity already differed as a function of future test order, we directly compared decoding time courses for items that were later tested early versus late, separately for Item 1 and Item 2 in the fixed-order and random-order conditions. No significant difference clusters were observed in either encoding epoch, in either condition (Supplementary Figure S2).

**Figure S1.**
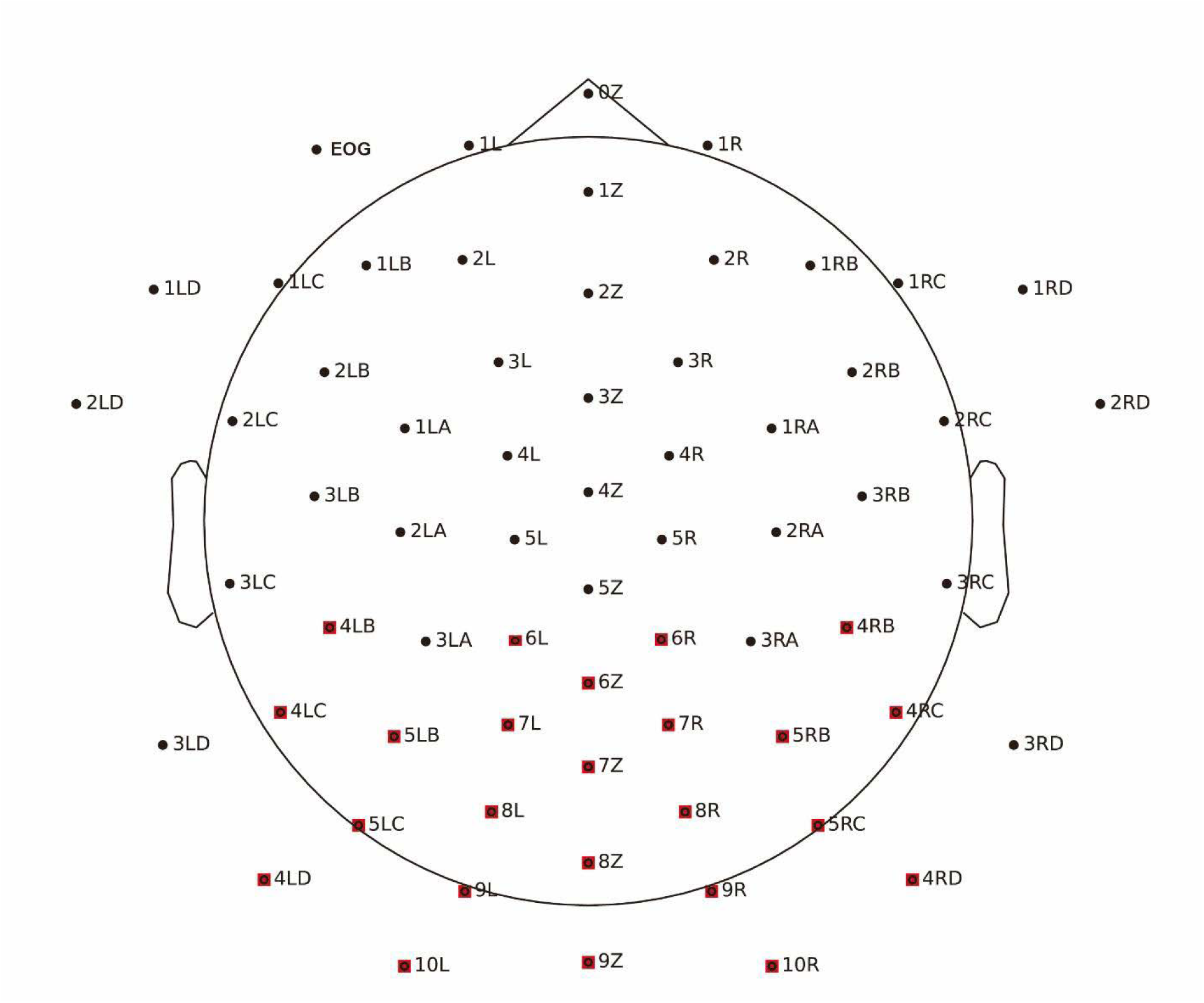
EEG cap layout. Two-dimensional topographical projection (top view) illustrating electrode positions relative to a representative scalp contour. The 24 most posterior channels selected for the decoding analyses are highlighted in red.

**Figure S2.**
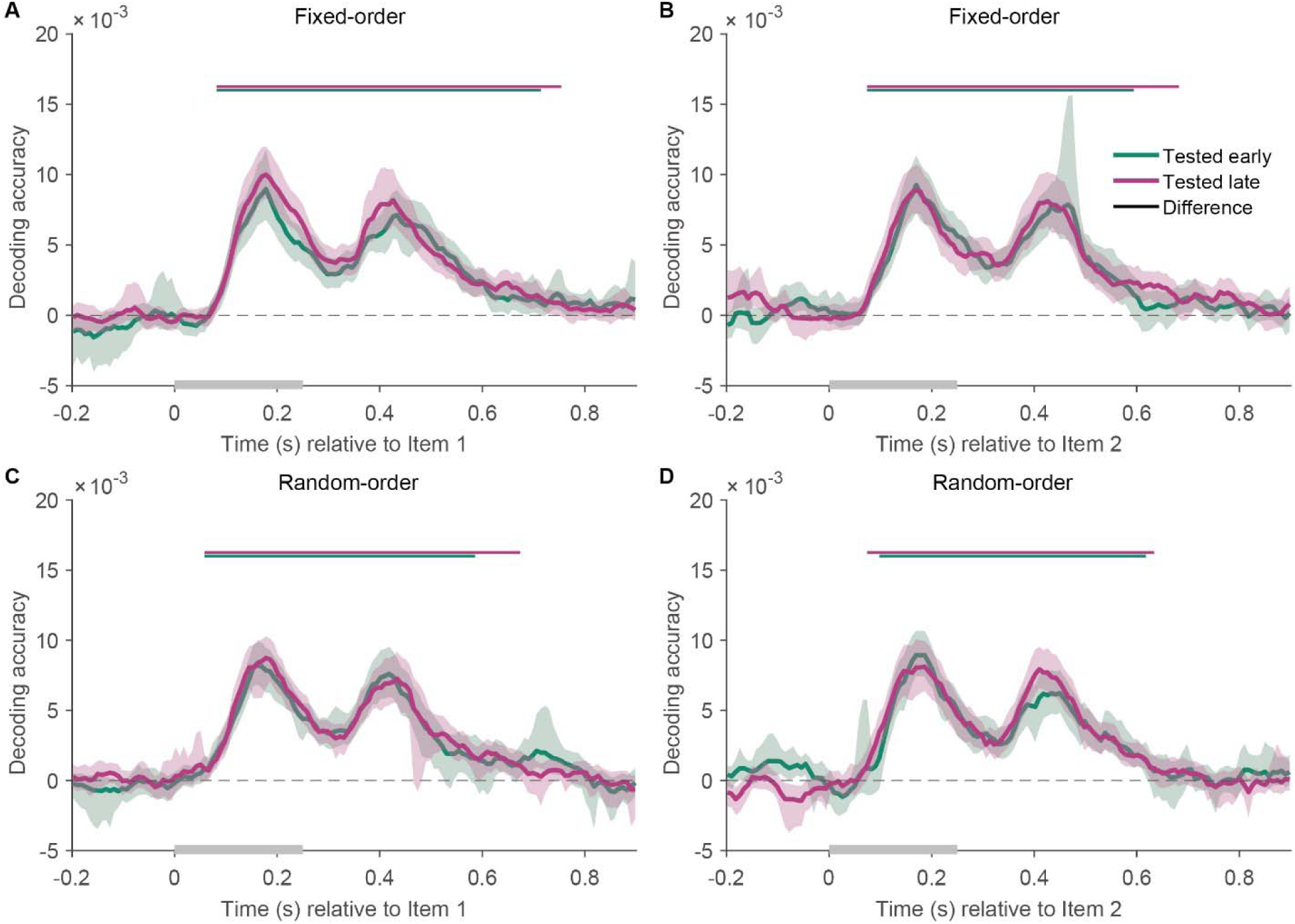
Future test order contrasts during encoding. Time-resolved voltage decoding during the encoding of Item 1 (A, C) and Item 2 (B, D), shown separately for fixed-order blocks (A, B) and random-order blocks (C, D). Within each panel, decoding traces are compared between items that were eventually tested early and items that were tested late. Black horizontal bars indicate significant differences between the two traces based on two-sided cluster-based permutation tests.

**Figure S3.**
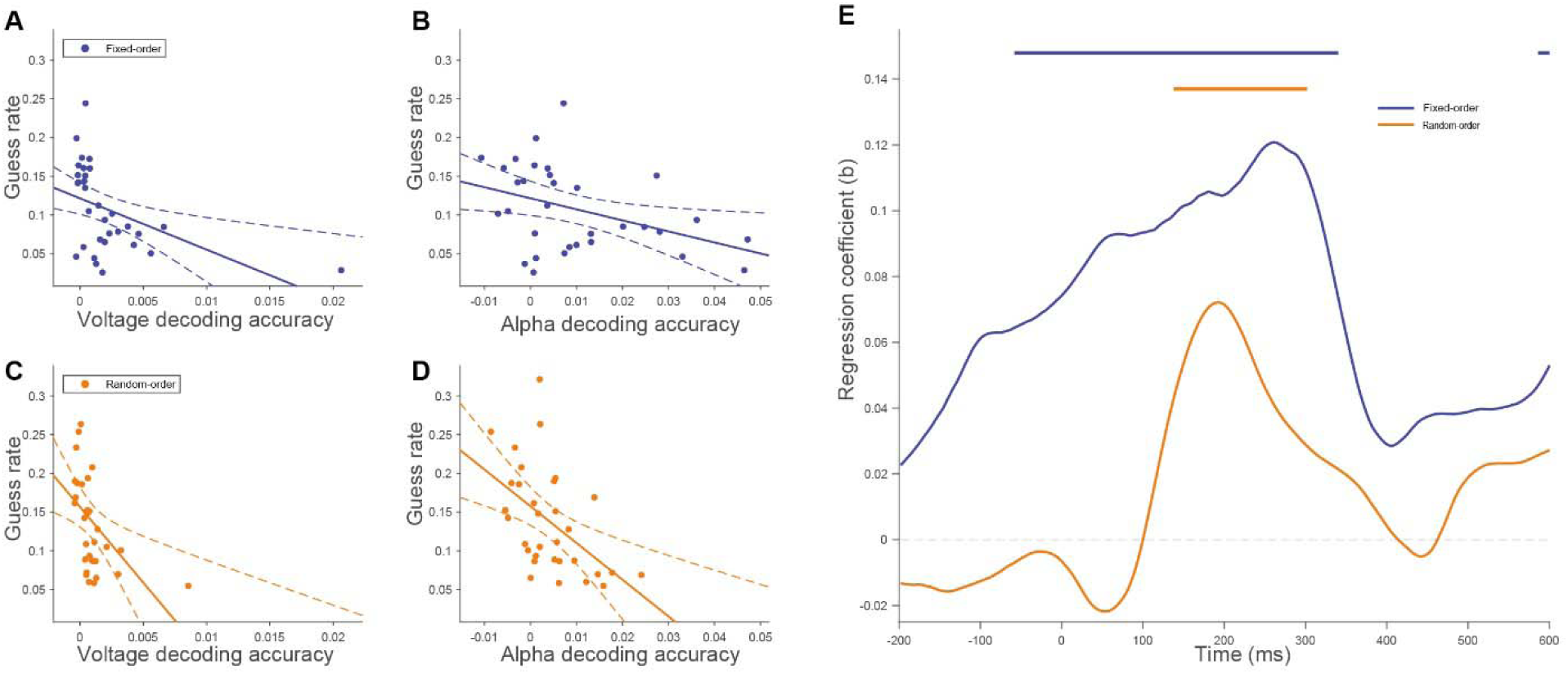
Regression analyses linking neural decoding to behavioral performance. (A–B) Fixed-order and (C–D) random-order associations of voltage and alpha power decoding with guess rate. (E) Time-resolved association between alpha power and voltage decoding. Horizontal bars indicate significant time points (p < 0.05).

## STAR□METHODS

### KEY RESOURCES TABLE

| REAGENT OR | SOURCE | IDENTIFIER |
| RESOURCE |  |  |
|---|---|---|
| Deposited data |  |  |
| Filtered and Epoched EEG Data | This paper |  |
| Behavioral Data | This paper |  |
| Software and algorithms |  |  |
| Customized MATLAB Scripts for data analysis | This paper |  |
| Customized R Scripts for data analysis | This paper |  |
| Palamedes Toolbox 1.8.2 | Nicolaas Prins and Frederick A. A. Kingdom | https://www.palamedestoolbox.org/ |
| EEGLAB 2023.1 | Swartz Center for Computational Neuroscience, University of California San Diego | https://sccn.ucsd.edu/eeglab/ |
| CircStat 2012a（1.21.0.0） | Philipp Berens | https://www.jstatsoft.org/article/view/v031i10 |
| R 4.3.3 | The R Foundation for Statistical Computing | https://www.r-project.org/ |
| MATLAB R2023b | The MathWorks, Inc. | https://www.mathworks.com/products/matlab.html |

### EXPERIMENTAL MODEL AND STUDY PARTICIPANT DETAILS

#### Participants

Thirty-two young adults (20 women; mean age 21.78 years, range 18–29) participated in the experiment. They were recruited via the University of Groningen first-year participant pool and received either course credit or monetary compensation. Participant selection relied on the successful completion of a pre-screening test, which was a shortened version of the main experiment (120 trials). The cut-off criterion for selection was an average accuracy above 70%. The sample size was based on earlier studies with similar designs, such as the one by Wolff et al.23. All participants were informed about the experimental procedures and the data sharing procedures. Informed consent was obtained electronically; only those who agreed were allowed to proceed with the experiment. The study was approved by the Ethics Committee of the Faculty of Behavioural and Social Sciences, University of Groningen (PSY-2425-S-0198), in accordance with the NETHICS code.

#### Apparatus and stimuli

The experimental stimuli were generated and controlled by Psychophysics Toolbox, a freely available Matlab extension^44^. The stimuli were presented on a 27-inch LCD screen running at 100 Hz and a resolution of 1920 by 1080 pixels. Viewing distance was approximately 65 cm, constituting a 36.49 px/° viewing angle. Participant responses were given via a two-button response box, with a white button on the left and a red button on the right. Ambient room lighting was maintained at approximately 100 lumen. Throughout the experiment, a gray background was displayed and all stimuli appeared at the center of the screen.

The fixation dot was black, 0.41° in diameter. Memory items and probes were sine-wave gratings with 20% contrast, 8.22° diameter, and a spatial frequency of 0.98 cycles/°. The phase of each grating was independently randomized for every presentation, both within and across trials. Memory item orientations were selected from six evenly spaced angles (4°, 34°, 64°, 94°, 124°, 154°). The two memory orientations were always different within a trial, and all possible orientation combinations occurred an equal number of times across the experiment. The impulse stimulus was a white disk (10.67° in diameter). Probe gratings were rotated by ±10°, ±25°, or ±40° relative to the corresponding memory orientations. Each probe contained a gray circular marker (1.23° diameter) at its center, in which a digit was displayed (“1” or “2”), referencing the memory item being probed. Feedback consisted of a white “O” for correct, a black “X” for error, or “Miss” for no response, shown to the left of fixation for Item 1 and to the right for Item 2 (independent of testing order).

### METHOD DETAILS

#### Procedure

The trial sequence is illustrated in Figure 1A. Each trial additionally included an initial fixation period (500-800 ms) and a trial-wise feedback display (300 ms), which are not depicted in the figure. Participants were required to memorize the orientations of Item 1 and Item 2, presented for 250 ms each with 900 ms between presentations. During each maintenance delay, only the fixation dot was on screen, with Impulse stimuli of 100 ms duration, appearing 500 ms before Probe onset. Upon presentation of the probe, participants judged whether the memory item indicated by the numerical cue would need to be rotated clockwise or counterclockwise to match the probe. Responses could be made within 1550 ms following probe onset, spanning the 250 ms probe display and the subsequent 1300 ms blank interval. The same response window applied to both probes. Responses were collected using an external response box, which featured a white button on the left and a red button on the right. Counterclockwise rotations were reported by pressing the left (white) button, and clockwise rotations by pressing the right (red) button.

The experiment consisted of 1680 trials, divided into four consecutive sessions that were separated by breaks. Participants could determine the duration of the breaks between sessions. Each session consisted of 14 blocks of 30 trials each, with an average performance score presented as feedback after each block. Prioritization was manipulated via a blocked design by informing participants of the upcoming test order. At the beginning of each experimental block, an on-screen text cue informed participants about the upcoming probe order. In the fixed-order condition, the message stated either “You will always be tested on Item 1 [or 2] first”, such that participants knew in advance which memory item would be tested earlier throughout the upcoming block. In the random-order condition, the message stated “The test order will be random.” In this condition, testing order was unpredictable on a trial-by-trial basis, yet balanced within each block. Within each session, all blocks followed the same probe order condition. The different probe order conditions were fully counterbalanced across participants using a Latin square design, and the six probe rotations were equally represented across the experiment.

For clarity, we provide a short summary of the terminology used throughout. *Memory items* refer to the stimuli in their encoding sequence, distinguishing between Item 1 (first presented) and Item 2 (second presented). *Impulse* refers to the task-irrelevant visual stimulus presented during the delay periods to probe quiescent neural representations. Impulse 1 appears during the first maintenance interval, and Impulse 2 appears during the second interval. *Probe* refers to the specific temporal instance of the retrieval event, denoted as Probe 1 (the first recall opportunity) and Probe 2 (the second recall opportunity). The *Tested Item* classifies memory items based on their testing order: Items recalled at Probe 1 are defined as items tested early, whereas those recalled at Probe 2 are items tested late. Finally, the *Prioritization* condition refers to whether this test order was known in advance (fixed order) or remained unknown (random order) prior to the retrieval phase.

#### Behavioral modeling

Memory performance was analyzed similarly to Wolff et al.^23^, using the Palamedes Matlab toolbox (http://www.palamedestoolbox.org/). A cumulative Gaussian psychometric function incorporating guess- and lapse rate terms was fitted to each participant’s proportion of clockwise responses across the six probe rotations, using maximum likelihood estimation. For each participant, data from both prioritization conditions were jointly fitted. Thresholds were constrained to be equal, while slopes, lapse rates, and guess rates varied freely. Modeling was performed separately for the item tested early and the item tested late, because these item classes differed in their priority status within the trial and were therefore expected to show distinct effects of prioritization. Within each joint fit, Bootstrap confidence intervals (CI) for the observed response proportions in each condition were obtained using 50,000 bootstrap samples.

#### EEG acquisition

EEG data were recorded using EEGGO software and a TMSI Refa 8–64 amplifier with 64 sintered Ag/AgCl electrodes arranged according to the equidistant system (see Supplementary Figure S1 for the electrode layout). The sampling rate was 1000 Hz, and electrode 5Z, located at the center of the cap, was used as the online reference. The impedance of all electrodes was kept below 5 k.

#### EEG preprocessing

The recorded data was re-referenced to both mastoids, bandpass filtered (0.1 Hz high-pass and 40 Hz low-pass), and down-sampled to 500 Hz, using EEGLAB. Independent Component Analysis (ICA) was first performed. Subsequently, the ICLabel plugin was used to identify and remove components associated with ocular artifacts, including eye blinks and eye movements. EEG data were then segmented into relatively long epochs from -500 ms to 1200 ms relative to the onset of the four stimuli (Items 1 and 2, and Impulses 1 and 2). This extended epoch length provided sufficient boundary padding for subsequent analyses on shorter analysis windows, particularly for stable alpha power estimation. Baseline correction was then applied using the pre-stimulus interval from -200 ms to 0 ms.

For the subsequent analyses, 24 posterior channels were selected (6L, 6Z, 6R, 7L, 7Z, 7R, 8L, 8Z, 8R, 9L, 9Z, 9R, 10L, 10R, 5LB, 5LC, 5RB, 5RC, 4LB, 4LC, 4RB, 4RC, 4RD, and 4LD). This posterior channel set was selected a priori based on the impulse based decoding approach of Wolff et al ^23^. Epochs were flagged for rejection using EEGLAB’s joint-probability procedure across non-EOG channels. Specifically, an epoch was marked when the normalized joint probability of its voltage values exceeded 5 standard deviations either for any individual channel or across channels considered jointly^45,46^.

#### Voltage decoding

Voltage decoding followed the methods described in Wolff et al., 2017 and 2020^23,47^. Item 1, Item 2, Impulse 1, and Impulse 2 epoch were defined as -200 to 900 ms relative to their respective onsets for subsequent decoding. However, to focus on the immediate neural response elicited by the impulse and avoid potential interference from the subsequent probe, statistical analyses and visualization for the impulse epoch were restricted to the -200 to 600 ms window. For Item 1 and Item 2 encoding epochs, decoding was performed specifically on the orientation of the stimulus presented during that interval. Within each epoch, trials were partitioned into four subgroups based on the combination of prioritization condition (fixed-order vs. random-order) and testing order (item tested early vs. item tested late). For Impulse 1 and Impulse 2 maintenance epoch, trials were split by prioritization condition. Within each split, decoding was performed twice: once using orientation labels corresponding to the item tested early, and once for the item tested late.

We decoded voltage data using a sliding window approach that takes into account the relative voltage changes within a 100 ms window^47^. Time points within 100 ms of each channel and trial were first downsampled by taking the average every 10 ms, resulting in 10 voltage values for each channel. Next, a dynamic baseline was applied by subtracting the mean activity within that same 100 ms window from each of these 10 values for each channel. All 10 voltage values per channel were then used as features for the five-fold cross validation decoding.

We decoded multivariate data based on Mahalanobis distance. For each fold, a covariance matrix was computed from the training trials using a shrinkage estimator^48^. This was used to determine the Mahalanobis distance between each test trial and the average multivariate pattern of all six presented orientations. To enhance the reliability of distance estimates, this procedure was repeated 50 times. In each repetition, the data were randomly partitioned into five folds via stratified sampling, and the number of trials for each orientation in the four training folds was equalized by randomly sub-sampling to the minimum count per orientation.

For each test trial, the six computed distances were centered with respect to the current orientation, then sign-reversed such that higher values indicate greater pattern similarity, to facilitate interpretation. Orientation decoding performance was defined as the centered distances multiplied by the cosine of the relative angles^23^, where higher values reflect stronger evidence for orientation-specific neural representations. The entire pipeline was applied in a time-resolved manner, sliding in 8 ms steps.

#### Alpha power decoding

We also used alpha power to decode the item specific information. To derive alpha power, we first applied a bandpass filter to the preprocessed EEG signal, isolating the 8–12 Hz alpha frequency band. Subsequently, a Hilbert transform was employed to compute the instantaneous amplitude of the filtered signal, representing the amplitude envelope of alpha power.

Alpha power decoding was conducted on impulse epochs only, using a -200 to 600 ms window relative to impulse onset. This choice was motivated by recent evidence suggesting that item specific information decodable from alpha power primarily reflects the attentional prioritization of information already held in WM, rather than a generic maintenance signal or a stimulus-locked encoding response^24^. To enhance the signal-to-noise ratio for alpha power decoding, pseudotrials were generated ^49^. Initially, the data, comprising six orientation categories, were subjected to random stratified sampling to partition them into five folds, ensuring an equal distribution of orientations across each fold. Trials within the same orientation were averaged within each fold, resulting in one pseudotrial per orientation category per fold. For cross validation, each iteration employed the pseudotrials from four folds as the training set, while the pseudotrials from the remaining fold served as the test set.

The resulting pseudotrials were then submitted to the same cross validated decoding framework as in the voltage analysis, and decoding accuracy was averaged across all repetitions. Unlike voltage decoding, however, no dynamic baseline was applied before alpha power decoding. Alpha amplitude was estimated from the Hilbert-derived envelope, which reflects the time-varying magnitude of alpha band activity^50^.

### QUANTIFICATION AND STATISTICAL ANALYSIS

Behavioral performance was quantified using the parameters derived from psychometric function fitting. Separate 2 2 repeated-measures ANOVAs were conducted on the slopes (representing precision) and guess rates, with prioritization condition and tested item as within-subject factors.

To evaluate the time course of orientation-specific neural representations for both voltage and alpha power decoding, cluster based, sign-permutation tests with cluster-mass correction were conducted within the Memory item epoch (0–900 ms) and Impulse epoch (0–600 ms). Each test was based on 100,000 permutations, using a sample-level cluster threshold of *p* < 0.05. We defined cluster mass as the sum of point-wise *t values* within each cluster. For tests against chance level within a single condition, one-sided permutation tests were employed, whereas for comparisons between two conditions, two-sided permutation tests were conducted. The reported *p values* reflect cluster level statistics obtained from this procedure.

For direct condition comparisons, we averaged the decoding accuracy within the Impulse 1 and Impulse 2 epoch. Specifically, we focused on the interval of 100–600 ms from impulse to probe onset. The mean decoding accuracy within these windows was subjected to a two-way repeated-measures ANOVA, with Impulse (Impulse 1 vs. Impulse 2) and prioritization condition as within-subject factors, to examine the effect of prioritization on neural representations over time.

To assess whether these neural decoding measures account for the behavioral prioritization effects, we examined the relationship between neural decoding and guess rate. For each participant and each prioritization condition, we separately averaged voltage decoding over the 100–400 ms post-impulse interval and alpha power decoding over the −200–600 ms impulse locked interval. The analysis windows were selected based on previous findings on voltage decoding^23^ and alpha power decoding^24^, as voltage decoding is mainly observed after impulse onset, whereas alpha power decoding can be observed before impulse onset and persists over a longer period. Specifically, this included decoding scores for the item tested early following Impulse 1 and for the item tested late following Impulse 2 in both the fixed-order and random-order conditions. Guess rate estimates were processed in the same way, yielding one overall guess rate per participant in each prioritization condition. We then compared three regression models within each condition to predict guess rate: a model with voltage decoding as the predictor, a model with alpha power decoding as the predictor, and a model including both predictors. To examine the temporal association between alpha power decoding and voltage decoding, we fitted separate linear regression models across participants at each time point from −200 to 600 ms, with voltage decoding averaged over 100–400 ms as the dependent variable and alpha power decoding at each time point as the predictor. This analysis was conducted separately for the fixed-order and random-order conditions.

To examine whether internal attention reduces WM guess rate by enhancing memory item representations and making them more accessible, we conducted mediation analyses with alpha power decoding as an index of internal attention and voltage decoding as an index of memory item representation, separately for the fixed-order and random-order conditions. Based on the temporal association between alpha power decoding and voltage decoding, alpha power decoding was examined during the pre-impulse (−200–0 ms) and post-impulse (100–400 ms) intervals, while voltage decoding was averaged over 100–400 ms. The mediation analyses were conducted both by combining Impulse 1 and Impulse 2 and separately for each impulse within each prioritization condition.

